# Live and inanimate predator-associated cues suppress the population of sap-feeding prey and induce polyphenism

**DOI:** 10.1101/2019.12.27.889634

**Authors:** Mouhammad Shadi Khudr, Tabea Dobberke, Oksana Y. Buzhdygan, Susanne Wurst

## Abstract

Non-consumptive effect of predation is a well-researched subject of which certain non-consumptive and predator-mimetic facets are yet to be investigated in plant-parasite systems

One clone of the green peach aphid *Myzus persicae* (Sulzer), raised on a model crop *Brassica oleracea* (L.), was exposed to different regimes of risks associated with ladybird *Coccinella septempunctata* (L.). This encompassed consumption, consumption alternated by non-consumptive effects, isolated predators, dead predator, predator dummy, as well as dummy, plants or soil cued with predator-borne suspension, and predator removal (exposure to plants previously visited and marked by a predator).

Over time, the respective risk regimes variably negatively impacted the prey population; the corpses, cued plants and dummies had considerable persistent negative effects on aphid reproductive success, contrary to the observation under predator removal. By the end of the experiment, polyphenism (winged morph production) also differed and was animated under the presence of a starved isolated predator; but faded when a predator corpse was present; and vanished under the dummy. Our findings, in this model aphid-crop system, contribute to the rapidly developing area of the ecology of fear, as we provide insights and novel means for aphid management that merit further examination across different eco-agricultural contexts.

## Introduction

Across natural and agricultural ecosystems, the impact of predation on the population dynamics and survival of prey is multifaceted. It takes place in the form of direct consumption and also occurs non-consumptively via fear of predation or intimidation (Lima 1998; Preisser et al. 2005; McCauley et al. 2011; Zöttl et al. 2013; Kersch-Becker and Thaler 2015). This can be in effect through different types of stimulation, such as the presence of impaired predators (i.e., predators altered experimentally to no longer consume the prey) (Nelson et al. 2004; Nelson and Rosenheim 2006), mere predator presence (McCauley et al. 2011; Kersch-Becker and Thaler 2015), and predation risk including induction by predator-borne cues (Preisser and Bolnick 2008; Ferrari et al. 2010; Ninkovic et al. 2013; Khudr et al. 2017).

### Information cues in action

The prey assess multiple cues and info-chemicals from their surroundings, including stimuli associated with their natural enemies (Lima and Dill 1990; Kats and Dill 1998; Lima and Steury 2005; Ferrari et al. 2010). Inducible prey defences after perception of environmental risk have been reported to be changeable relevant to the risk perceived and hence can be adjustable and adaptive (e.g., through transgenerational effects in aphids) (Keiser and Mondor 2013), as they may lead to fitness gains (Evans and Schmidt 1990; Coslovsky and Richner 2011; Zöttl et al. 2013). Responses to predation risk may span feeding cessation, escape and avoidance of predators (Nelson 2007; Keiser and Mondor 2013), dispersion (Roitberg et al. 1979, Nelson and Rosenheim 2006, Hatano et al. 2010), and habituation to risk (Shalter 1984; Holomuzki and Hatchett 1994). Also, prey reproductive success may undergo inhibition, or reproduction may be altered under intimidation by predators (Preisser et al. 2005; McCauley et al. 2011), alarm pheromone effects (de Vos et al. 2010), and due to behavioural changes (Nelson et al. 2004; Hoki et al. 2014). The effects brought about by predator cues (Norin 2009) diffuse through the aphid population and intensify because of communication by way of pheromones (de Vos et al. 2010; Keiser 2012, Ingerslew and Finke 2017), thereby inducing forms of phenotypic plasticity such as alata production (Weisser et al. 1999), or changes in reproductive success, or both (Khudr et al. 2017). Whichever choice the prey make, the decision of developing anti-predator responses and reacting to environmental risks incur an ecological cost (Agabiti et al. 2016; Hermann and Landis 2017; Ingerslew and Finke 2017), leading generally to compromised prey fitness and altered dynamics and behaviours (Lima and Dill 1990; Sih 1994; Sih 1997; Nelson et al. 2004; Preisser et al. 2005) to the advantage of survival (Francke et al. 2008; Keiser 2012). Using isolated predators or inanimate objects cued by predator info-chemicals as a form of risk to affect parthenogenetic phloem-feeding insects has not received enough attention compared to the effects of the employment of impaired predators (Nelson et al. 2004; Nelson and Rosenheim 2006). The lingering question remains, however, on what sensory modalities are the most important in affecting prey perception of predation risk and shaping prey response to the risk over time.

### Brevity versus longevity of non-consumptive effects

The evidence for the wide-spread impact of fear of predation is mounting across a vast array of taxa (Preisser et al. 2005; Hermann and Landis 2017). However, the current understanding of the role of the duration of risk exposure on how non-consumptive effects influence the biology of phloem-feeding insects is meagre, but see (Van Dievel et al. 2016). Prolonged or more frequent exposure to high frequencies/intensities of risk imposed by natural enemies can cause a state of environmental uncertainty for the prey (Sih 1992; Koops 2004; Trussell et al. 2011) that may dilute the strength of non-consumptive effects over time. This may alter the assessment of risk by the prey and concomitant prey responses against risk cues and thus may eventually induce prey habituation to risk, which can be contextual and contingent on prey physiology and energy dynamics (Trussell et al. 2011; Matassa and Trussell 2014). Moreover, unlike older predator-borne cues, fresh ones might be associated with higher localised risk and could then provide more certainty/reliability resulting in faster and more adaptive prey responses (Dixon and Agarwala 1999; Podjasek et al. 2005; Keiser and Mondor 2013) which could reduce ecological costs (Koops 2004; Ninkovic et al. 2013; Zöttl et al. 2013). Nevertheless, temporal aspects of anti-predator responses remain obscure, including whether and how constant versus fluctuating risks may affect prey population dynamics in the short and the long term (Lima and Bednekoff 1999; Hamilton and Heithaus 2001).

The intensity and consistency of the effects of predator-borne cues, when consumption and/or non-consumptive effects change through time, may result in trait-mediated prey responses; the effect of which may scale up through ecosystems (Lima and Bednekoff 1999, Steffan and Snyder 2010; Matassa and Trussell 2014). To date, comprehensive studies on profiling reproductive success and phenotypic plasticity of parthenogenetic phloem-feeding prey under respective exposure to an array of non-consumptive risk types over a given period are scarce. Although prey intimidation by the constant presence of an isolated predator (non-consumptive effects) has been demonstrated before in insects (e.g., dragonfly) (McCauley et al. 2011), this has never been sufficiently tested over a generation time in the presence of isolated predators or especially when different types of inanimate predator-imitating cues are applied in model crop-aphid systems.

### Inducible defences and non-consumptive effects via biomimicry

Under favourable mesic conditions, the population of parthenogenetic phloem-feeding aphids comprises female congeners with very strong maternal/transgenerational effects (Mousseau and Fox 1998; Weisser et al. 1999; Hu et al. 2018). Responses to environmental stimuli become escalated by maternal preconditioning of daughters and granddaughters by means of telescoping generations (Kindlmann and Dixon 1989), as consecutive generations of offspring may develop and become preconditioned simultaneously (Mousseau and Fox 1998; Weisser et al. 1999; Hu et al. 2018), leading for instance to the production of winged aphids (alates, dispersive morphs) (Weisser et al. 1999; Kunert et al. 2005). The latter is an anti-predator trait of phenotypic plasticity; a polyphenic response to predation risk (Dixon and Agarwala 1999; Mondor et al. 2005); alates disperse to settle aside from the adverse circumstances that induced their generation (Dixon and Agarwala 1999; Kunert and Weisser 2003; Mondor et al. 2005).

Ladybirds are effective in aphid biocontrol (Hoki et al. 2014), particularly in greenhouse settings (Riddick 2017), and their footprints are known to elicit avoidance behaviour in aphids (Ninkovic et al. 2013). Aphids, mainly chemically detect the presence of ladybirds (e.g., *Coccinella septempunctata*) through exposure to the tracks and excretions left by the predator (Ninkovic et al. 2013), with evidence for sex-dependency as well as concentration effects of ladybird olfactory cues on aphid response (Youren 2012). Roitberg et al. (1979) reported that coccinellid predation risk induced aphids to scatter on plants and in between plants, which was followed by reduced reproductive success, indicating energy costs, allocated for the preconditioning of progeny (escapees), at the expense of reproduction (Mackay and Wellington 1975; Roitberg et al. 1979; Keiser and Mondor 2013). Furthermore, Fievet et al. (2008) demonstrated that exposure to indirect cues in the form of dead conspecifics due to predation induced behavioural changes that were accompanied by a decline in the aphid population. A relation between reduced feeding time and dwindling reproductive success (Sih 1992) has been described by Nelson (2007), as well. The reduction of feeding time was related, in the cited work, to predator disturbance frequency, where the presence of non-consumptive (impaired) predators led to reduced aphid fitness (Nelson et al. 2004). Optimal habitats for prey may change with time, as do anti-predator behaviours, thereby forcing the prey to resort to safer micro-habitats. However, avoidance of ecological risk can become increasingly costly on poor or scarce resources (Turner 1997), where optimal feeding sites are rare; changes in aphid reproduction may have variable extended effects on other trophic levels, highlighting the far-reaching effects of fear of predation over time (Preisser et al. 2005; Creel and Christianson 2008; Steffan and Snyder 2010).

It has been well-established that aphids respond to non-consumptive cues associated with their predators. For example, impaired predators have been shown to negatively impact aphid reproduction (Nelson and Rosenheim 2006). Encounters between green peach aphid *Myzus persicae* and corpses of its lacewing predator *Chrysoperla carnea* (Stephens) or plants pre-treated with lacewing cues have also been reported to induce anti-predator avoidance behaviour and lead to a differential reduction in aphid fitness as a consequence of fear of predation (Khudr et al. 2017). The literature provides examples of research on the employment of predator replicas (false predators) (e.g., Mitrovich and Cotroneo 2006) or artificial prey (e.g., Morrell and Turner 1970; Roslin et al. 2017) to study the dynamics of predator-prey and risk-prey interactions, but these studies usually focus on higher taxa rather than insects. As such, mimicking natural enemies by exposing aphids to cued and non-cued predator simulacra highly resembling the enemies has not been done before. Defining the effects of these different classes of non-consumptive cues to test whether their mimicry effects can repress aphid reproduction and induce concomitant anti-predator defences of extreme phenotypic plasticity deserves careful examination. For that matter, while most research on the ecology of fear focuses on changes in prey performance, following behaviour modifications, there is an emergent need to decipher previously uncharted areas of non-consumptive effects of predation risk on prey fitness and pest control where innovative ecological means are employed (Ingerslew and Finke 2017).

In the present work, we investigate the effects of exposure to various types of consumptive and non-consumptive risks, associated with and/or mimicking an aphidophagous insect *C. septempunctata* (L.), on the reproductive success (total aphid numbers per census) and extreme phenotypic plasticity (polyphenism) by counting winged aphid morphs at the end of the experiment. We use one genotype of a significant crop pest that is green peach aphid *M. persicae* (Sulzer). The rationale of the treatment setting was mainly built to draw on the developing area of knowledge on aphid responses to environmental cues and change based on their multi-sensory perception (visual, olfactory, chemical, tactile, vibrational, and via interoception) (Döring and Chittka 2007; Ninkovic et al. 2013; Barron and Klein 2016, Klein and Barron 2016, Khudr et al. 2017, Cabej et al. 2019; Tamai and Choh 2019). See (Fig. 1) for an overview of the conceptual design of the work.

**Fig. 1.**
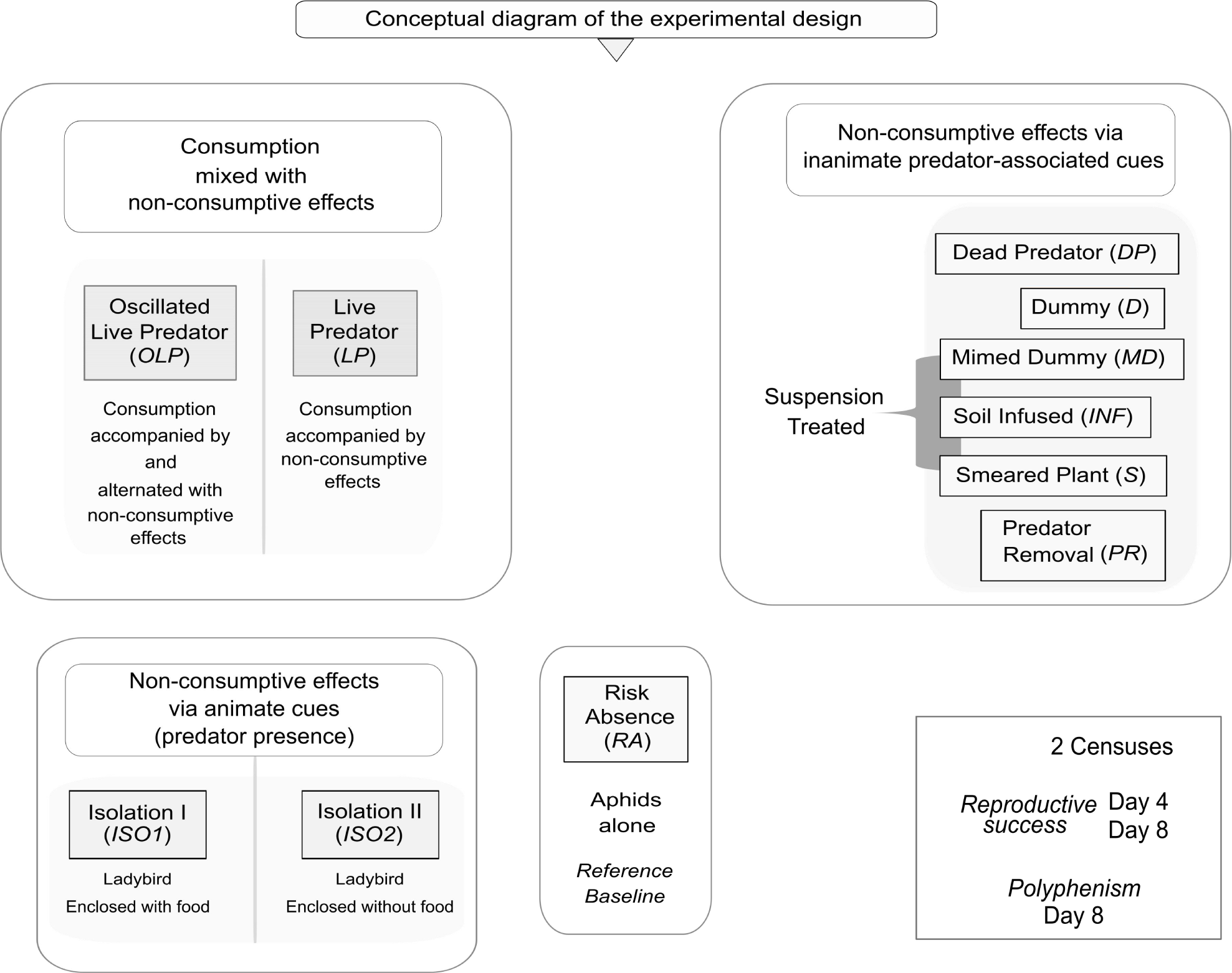
Schematic diagram of the conceptual set-up of the experiment. The diagram illustrates the characteristics of the applied risk treatments associated with ladybird *C. septempunctata* and the structure of the analysis of total numbers (as a proxy for reproductive success) on Day 4 and Day 8, and winged morphs (alata production denoting distinct phenotypic plasticity [polyphenism]) on Day 8 of green peach aphid *M. persicae* under the effects of the risk treatments.

### We test the following predictions

1. Dependent on the risk type and its longevity, different types of predator-associated risks are of similar magnitude in their repressive capacity of the target clonal aphid population, with consistent induced levels of phenotypic plasticity (marking escape tactics).
2. Aphid clonal population reacts almost the same to constant and oscillated presence of a predator.
3. The impact of an isolated predator, feeding on aphid conspecifics, is tantamount to the impact of the isolated predator deprived of the feeding.
4. Predator dummies would show a similar effect to that of the dead predator on the aphid population and that might not change when the dummies are enhanced with predator-borne cues.

## Materials and methods

### Organisms

This small-scale ecological experiment was performed at the Freie Universität Berlin, Germany, greenhouse conditions (22-24°C and 16:8 L:D). Savoy cabbage *Brassica oleracea var. sabauda*, cultivar Vertus 2 of stable agronomic performance, purchased form (Sperli©, Germany), was the treatment host plant. Two seeds were sown on opposite sides of plastic potware (11cm height and diameter) filled with steam-sterilised soil. All the treatment plants were 30 days old on Day 0 of the experiment and watered as needed.

To control for aphid preconditioning, one clonal lineage of *M. persicae*, initially supplied by Julius Kühn-Institut (JKI), Berlin, was reared for several generations on kale *Brassica oleracea* var. Sabellica (L.), cultivar ‘Lerchenzunge’, seeds purchased from (Quedlinburger Saatgut© through a local supplier, Berlin, Germany), and maintained under the above-mentioned conditions prior to the commencement of the experiment. Culture and treatment plants are both attractive for *M. persicae*; an important polyphagous pest (Blackman and Eastop 2007). A population of adults (females) of *C. septempunctata* (seven-spotted ladybird) was obtained from a commercial supplier (Katz Biotech AG, Baruth, Germany) to control for the sex, age, and background variability of the predator. After receipt, ladybirds were kept in the fridge (9°C), for one night, in plastic containers that included paper strips. Before trials, the refrigerated ladybirds were fed individually with 5 nymphs of *M. persicae* from the stock culture. Additionally, 30 ladybirds (∼1,067g in total) were frozen for 48h and utilised afterwards to prepare predator suspension (described below). The freezing of ladybirds occurred once delivered from the supplier, with no food provided before freezing.

### The setting

The microcosm (enclosure) of the experiment was made by installing a grid to hold up an ultra-fine mesh fitted with a zip that was used to enclose each plant pair with their pot. Twenty apterous third-instar clonal female nymphs of *M. persicae* were placed per microcosm, midway of the two plants per microcosm. The aphids were carefully transmitted, into each microcosm, on a rectangle of waxed paper (2×1cm), using a fine wet brush. The assay was run for 8 days, starting with Day 0 as the set-up day. Aphids were counted halfway (Day 4, 1^st^ census) and on the last day of the experiment (Day 8, 2^nd^ census) to determine treatment effects on the total raw numbers of *M. persicae*; the leaf morphology and the seedling architecture of savoy cabbage, the sedentary nature of aphids, and the controllable detachable enclosure of the microcosm all facilitated a careful aphid count on Day 4 with the least perturbation possible. On the final count on Day 8, alata production (i.e., the number of alates [winged morphs indicating polyphenism as discrete of phenotypic plasticity]) was also surveyed per microcosm. There were 11 simultaneous treatments × 7 repeats (microcosms) per treatment × 20 aphid instars per microcosm totalling up to 1540 genetically identical aphids. The treatments were as follows:

#### 1 Risk Absence (RA)

Aphids were alone in the microcosm, risk-free.

For treatments (2-5), ladybirds were singly introduced into the microcosms following 1h tune-up off the fridge at 24°C in individual petri dishes in order to ensure an unfazed transition into the ambiance of the experiment.

#### 2 Live Predator (LP)

Once aphids were transferred into respective microcosms, one ladybird was introduced per microcosm and remained enclosed for the full experiment time where it was left free to forage the microcosm. For aphid counting on Day 4, ladybirds were captured momentarily in separate plastic containers. After aphid census, each ladybird was relocated into its corresponding microcosm. The risk effect here was mainly consumption, but also the predator roaming the microcosm could generate a non-consumptive effect by their bio-signature cues (e.g., semiochemicals, tracks and excretion).

#### 3 Oscillated Live Predator (OLP)

The predator was left in the microcosm for 2 days followed by one day ‘out of microcosm’ and so forth periodically until the end of the experiment. We kept the ladybirds in the fridge (9°C), during the ‘time-out’, in separate glass containers covered with mesh and one provision of 5 stock of *M. persicae* nymphs as a feed. Containers were numbered and every ladybird was returned to its original corresponding microcosm after 1 h transitional period in individual petri dishes at 24°C. For the count on Day 4, ladybirds were captured momentarily in separate plastic containers. The risk effect here entailed phases of (ladybird in microcosm) including periodic consumption accompanied by possible non-consumptive effect due to microcosm-roaming by the predator, which was alternated with intimidation phases (intervals of ladybird out of microcosm) by inanimate ladybird-borne cues (e.g., semiochemicals, tracks and excretion) left within the environment of the aphid clone in the microcosm. This presence/absence rhythm of predation/predation risk served the purpose of creating short-term intervals of fluctuation/oscillation in risk exposure.

Treatments (4-11) all exclude direct prey consumption, and focus on prey intimidation resulting from the influence of different *animate* or *inanimate* cues associated with non-consumptive effects of the coccinellid predator; in these treatments the cues were attached to one plant only of the two available in the microcosm, referred to as the ‘cue-treated plant’.

#### 4 Isolation I (ISO1)

Each ladybird was enveloped with 20 aphids to feed on, in a fine-mesh sachet (3×3cm). The mesh was firmly sealed with a rubber band and tied to the stem of one plant in the microcosm using a string (3cm above soil surface). Instantaneously, aphids were transferred into the microcosm. Ladybird individuals were left enveloped continuously throughout the treatment. The risk effect here was prey intimidation (non-consumptive) via the constant presence of an isolated animate predator, emitting semiochemicals, including olfactory cues, underlain by concurrent diffusion of alarm pheromone by the consumable conspecific aphids confined also in the predator sachet.

#### 5 Isolation II (ISO2)

Ladybirds were individually enveloped, as mentioned above in *ISO1*, but without the aphid feed. On Day 4, during aphid census, the ladybirds were captured briefly and fed with 5 aphids from the stock culture in separated plastic containers before they were re-enveloped in the corresponding sachets. The risk effect here was non-consumptive via the *mere* presence of an animate predator emitting semiochemicals, including olfactory cues, (without consumption of prey conspecifics in the predator sachet).

#### 6 Dead Predator (DP)

Fourteen ladybirds were frozen. Then, two randomly selected ones were transferred into each microcosm; one ladybird was placed on the ground close to the stem, while the other was carefully tied to the plant stem (3cm above soil surface), using a sewing kit. The risk effect here was non-consumptive by inanimate predator-borne cues (visual and olfactory/chemical). This treatment draws on our recent work on the effects of predator cadavers on aphid population growth (Khudr et al. 2017).

#### 7 Dummy (D)

Mock ladybird simulacra were prepared using shaped masses of odourless synthetic paste and colour markers; visually mimicking *C. septempunctata*. The dummies were waterproofed beforehand, using a transparent polish. The first dummy was placed on the soil surface, close to the stem. The second was tied to the plant stem (as in the treatment *Dead Predator*). A string was attached to the dummy’s end to tether it to the stem, (Fig. 2). The risk effect here was non-consumptive by inanimate cues (visual mimicry).

**Fig. 2.**
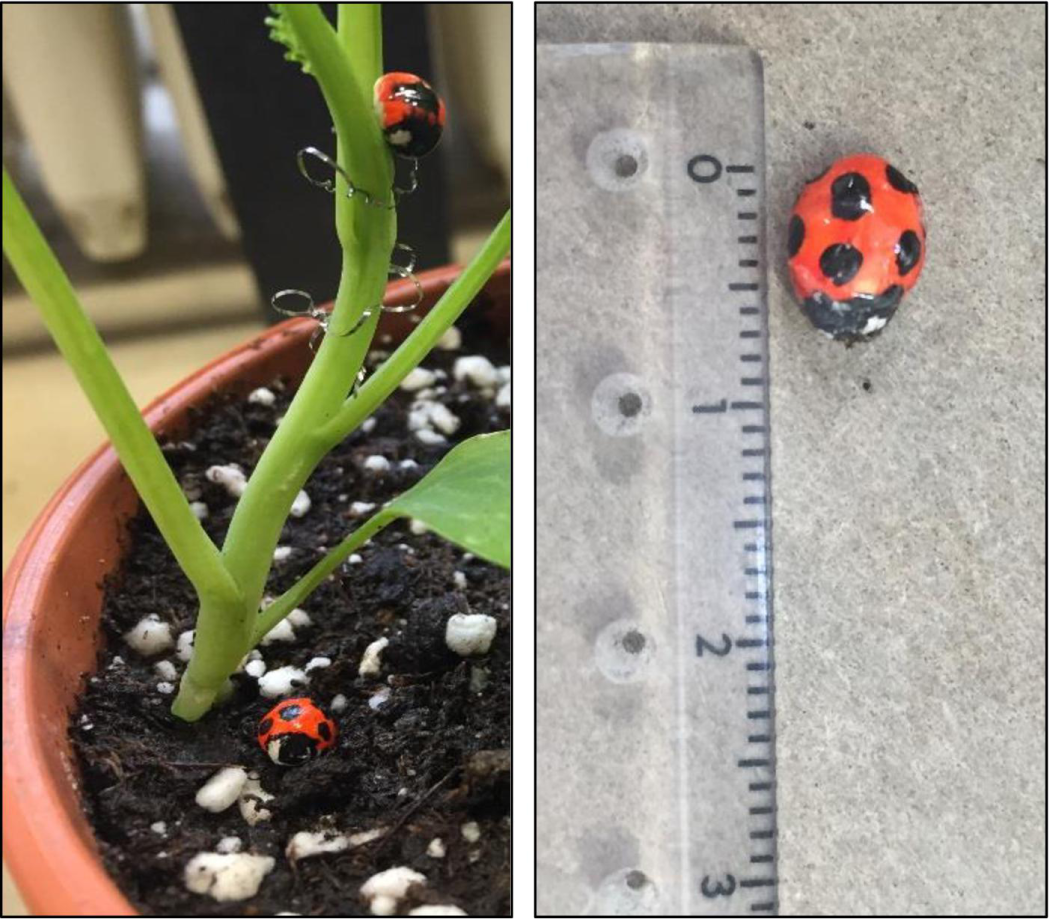
Predator dummy. The composite photograph displays the replica or simulacrum of ladybird *C. septempunctata* (termed as ‘dummy’). The same dummy was cued with the ladybird suspension, conveying predator-borne chemical cues, as described in the main text to create the decoy termed ‘mimed dummy’. The left part represents the set-up of the dummy or the mimed dummy, while the part on the right represents an exemplary dummy with reference in cm.

For the treatments (8-10), the above-mentioned predator suspension was prepared by fine-crushing 30 frozen ladybirds, using a grinding glass kit. Water was added up to a volume of 150ml. The suspension was vigorously shaken before each application.

#### 8 Mimed Dummy (MD)

Dummies were bound to one plant in the microcosm as described in the *Dummy* treatment above. Then, they were carefully smudged several times with the suspension prior to the introduction of aphids into microcosms. The risk effect here was non-consumptive by inanimate mimicry of the predator as the *MD* in this manner potentially provided aphids with two types of information: visual cues of the predator dummy and olfactory or chemical stimuli due to the added predator-borne cues to the surface of the dummy.

#### 9 Smeared Plant (S)

The leaves and stem of one plant in each corresponding microcosm were generously daubed with the suspension prior to the introduction of aphids. The risk effect here was non-consumptive by inanimate predator-borne info-chemicals.

#### 10 Soil Infused (INF)

A syringe with a modified elongated nozzle was inserted into the soil to gradually inject the aforementioned suspension at varying depths in contact with the roots of one plant of the two available in the microcosm. The infusion process of 10 mini-shots (1.9 ml per shot) covered the root system up to the disc where the stem came forth; 19 ml suspension was used in total per microcosm. This created a fine potential ‘prey intimidation zone’ in and on the soil surrounding the root system and the stem base of the treated plant. The risk effect was non-consumptive after the soil-infusion with predator-borne info-chemicals, drawing on our recent work on the applied infusion method for soil manipulation to directly or indirectly impede pest population growth (Khudr et al. 2017).

#### 11 Predator Removal (PR)

Ladybirds were taken off the fridge for 1h to tune-up and fed with 5 nymphs of *M. persicae* from the stock population. For 48 h, one ladybird was left with one plant only per microcosm for each repeat of this treatment in order to have the plants marked by the bio-signature of the ladybird. Subsequently, after the removal of the predator, a fresh plant (untreated and of the same age) was carefully transferred into each microcosm, with a mass of soil surrounding the roots. This was followed by the introduction of aphids. The risk effect here was non-consumptive by inanimate predator-borne cues (e.g., semiochemicals, tracks, excretions).

See (Fig. 1) for a schematic conceptual diagram of the factors/cues underlying the predator-associated risks, and (Supplementary Table S1) for further details.

### Statistical analysis

We used R version R 3.3.1 (R Core Team 2016). First, to examine the repressive effect of each predator-associated risk treatment on aphid total numbers per plant in the microcosm (a proxy for reproductive success) with two counts on Day 4 and Day 8, we applied a generalised linear mixed effect model (GLMM), with the function ‘glmer’, ‘bobyqa’ optimisation, and gamma family (due to the non-normal shape of distribution [positive skewness], and as confirmed by Shapiro test), using R package ‘*lme4*’ (Bates et al. 2015). The microcosm was nested within the count day times and randomised in the model; the main effects of the models were respectively revealed using an Anova command, R packages ‘car’ (Fox and Weisberg 2011). The predator-associated risk treatment comprised 11 levels (fixed effects) as specified above; the treatment “Risk Absence (*RA*)” made the model baseline. This was followed by a posthoc pair-wise multiple comparison test (Tukey’s HSD), R package ‘lsmeans’ (Lenth 2016).

Second, we investigated the effects of the aforementioned predator-associated risk treatments on alata production as alates were counted on Day 8 at the end of the experiment. We applied a generalised linear model (GLM) with a quasipoisson family (due to over-dispersion and non-normality [confirmed by Shapiro test]), using R package ‘multcomp’ (Hothorn et al. 2008). The effects were: 1) The 11-level risk treatment, where “Risk Absence (*RA*)” was the model baseline, 2) Aphid total numbers per microcosm (aphid density) as a covariate, and 3) the interaction between these effects. The main effects of the models were revealed using an Anova command, R packages ‘car’. See (Fig. 1) and (Supplementary Table S1) for conceptual and tabular schemas of the design. Data for this study are available at the figshare repository via the following URL: https://figshare.com/s/b99e89b8a3c6160ab74f

## Results

### Aphid reproductive success

As shown in (Fig. 3), aphids suffered a clear loss in reproductive success that was contextual and contingent upon the treatment (χ2_(10,262)_ = 69.07, P < 0.0001), underlain by the corresponding imposed predator-associated risks namely: constant mixed consumptive and possible non-consumptive risks (*LP*) (P < 0.0001), oscillated consumptive risk with intervals of exposure to non-consumptive risk associated with predator-borne cues (*OLP*) (P < 0.0001), non-consumptive animated risk associated with an isolated predator enclosed with an aphid conspecific feed (*ISO1*) (P = 0.019), non-consumptive animated risk associated with an isolated predator without the aphid feed (*ISO2*) (P = 0.011), non-consumptive risk associated with a predator corpse (*DP*) (P = 0.0003), non-consumptive risk associated with a predator replica (*D*) (P = 0.013), non-consumptive risk associated with a predator replica treated with predator-borne cues (*MD*) (P = 0.003), and non-consumptive risk associated with host plant treated with predator-borne cues (*S*) (P = 0.004).

**Fig. 3.**
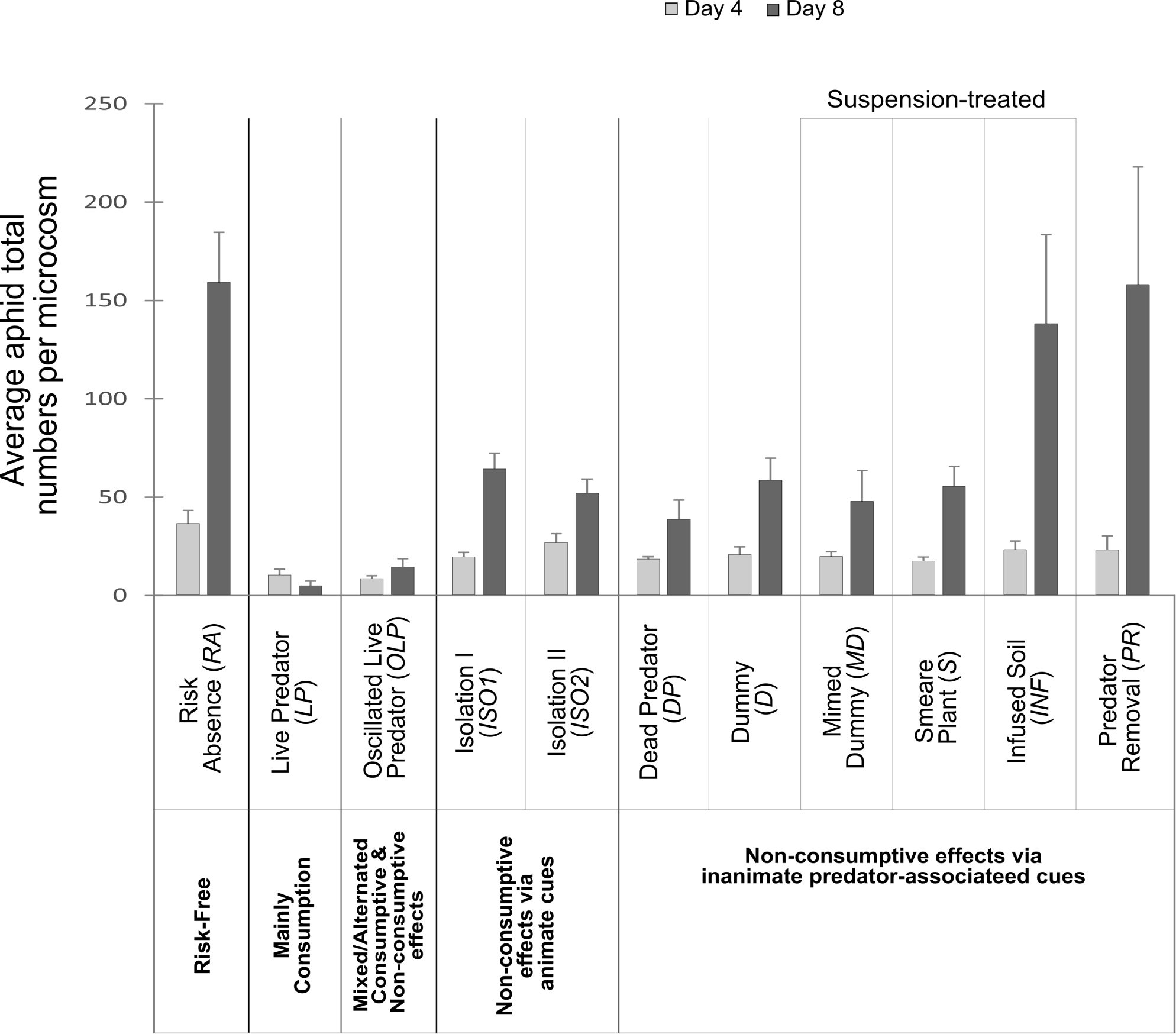
Aphid reproductive success under predator-associated risks. The chart illustrates the average aphid total raw numbers (±SEM) in the microcosm (as a proxy for the reproductive success of green peach aphid *M. persicae*) in response to the risk treatments associated with ladybird *C. septempunctata*. The light grey bar shows the mean reproductive success on Day 4 (1^st^ census); the darker bar displays the mean reproductive success on Day 8 (2^nd^ census).

The repression of the reproductive success was lower than in the control for all treatments, albeit being less noticeable under both the non-consumptive risk associated with soil-infused predator-borne cues in (*INF*) and the non-consumptive risk associated with host plant previously exposed to interaction with a live predator in (*PR*), as the effects of *INF* and *PR* were statistically insignificant (see Supplementary Tables S2 for model details, and Table S3 for posthoc multiple pairwise comparisons).

*LP* (x-bar = 10 ±3SEM aphids in the 1^st^ census [Day 4], and x-bar = 5 ±2SEM aphids in the 2^nd^ census [Day 8]) and *OLP* (x-bar = 8 ±3SEM aphids in the 1^st^ census, and x-bar = 14 ±4SEM aphids in the 2^nd^ census) had alternately together the strongest negative impacts on aphid numbers throughout the experiment, (Table 1, Fig. 3). The least effective treatment on Day 4, ranking 10^th^ in impeding aphid reproductive success, was (*ISO2*) (x-bar = 26.57 ±4.59SEM aphids) compared to the control (*RA*) (x-bar = 36 ±6.65SEM aphids). The least impeding one on Day 8 was (*PR*) with a very poor decrement of aphid reproductive success (x-bar =158 ±60SEM aphids). The treatments (*ISO1* on Day 4, and *ISO2* on Day 8) ranked 5^th^ on the impediment scale; whereas, the 3^rd^ and 4^th^ ranks were always held by (*S*) and (*DP*) on Day 4, and (*DP*) and (*MD*) on Day 8, as they were more efficient in their impeding effects than *ISO1* and *ISO2*, (Table 1, Fig. 3). It is noteworthy that (*S*) of the treatments excluding a live predator had the most negative impact on aphid reproductive success (x-bar = 17 ±2SEM aphids) on Day 4, that was 52% decrement of reproductive success compared to the observation under the control (*RA*); nevertheless, (*S*) was only 23% and 20% less effective than (*OLP*) and (*LP*), respectively. At the bottom of the chart was (*ISO2*) with only 25% reduction of reproductive success in comparison with the control, (Table 1, Fig. 3).

Noticeably, in the 2^nd^ census the risks associated with inanimate cues led to nuanced differential effects of impeding aphid reproductive success: *Dead Predator* (*DP*), being the most effective, resulted in ∼ 76% lower reproductive success (x-bar = 38 ±10SEM aphids) than the control (x-bar = 159 ±26SEM aphids); (*DP*) was only 21% and 15% less effective than (*LP*) and (*OLP*), respectively. By contrast, the aphid population thrived by the 2^nd^ census and reached a high score after the soil was cued in (*INF*) (x-bar = 138 ±45SEM aphids) *and also* under *Predator Removal* (*PR*) (x-bar = 158 ±60SEM aphids), entailing short-lived risk effects. Further, (*DP*) maintained the same rank (7^th^) in both censuses; likewise (*INF*) kept the 9^th^ rank across censuses. Additionally, (*DP, S*, and *MD*) were always within the rank range 3^rd^-6^th^, and (*D*) always maintained the 7^th^ rank across censuses (Table 1).

**Table 1.** Relative impediment of aphid reproductive success by predator-associated risks in ranks across two censuses. The different risk treatments are ordered in terms of the decrement of aphid numbers in the microcosm (as a proxy for reproductive success of green peach aphid *M. persicae*) in response to the risk treatments associated with ladybird *C. septempunctata*. Each treatment has an impediment rank number above and a percentage beneath that elucidates the decrement of the aphid population relative to the risk-free control (Risk absence [*RA*]). For example, on Day 8 (1^st^ census), aphid reproductive success under (*DP*) was 76% lower than the reproductive success under (*RA*), i.e., the reproductive success in (*DP*) was 24% the reproductive success under (*RA*). The treatments were: Risk Absence (*RA*), Live Predator (*LP*), Oscillated Live Predator (*OLP*), Isolation I (*ISO1*), Isolation II (*ISO2*), Dead Predator (*DP*), Mimed Dummy (*MD*), Dummy (*D*), Smeared Plant (S), Soil Infused (*INF*), and Predator Removal (*PR*).

| Rank | Most<br>efficient<br>① | ② | ③ | ④ | ⑤ | ⑥ | ⑦ | ⑧ | ⑨ | Least<br>efficient<br>⑩ | Risk<br>Free<br>⑪ |
| --- | --- | --- | --- | --- | --- | --- | --- | --- | --- | --- | --- |
| 1<sup>st</sup> Census<br>(Day 4) | *OLP*<br>77% | *LP*<br>72% | *S*<br>52% | *DP*<br>50% | *ISO1*<br>46% | *MD*<br>44% | *D*<br>42% | *PR*<br>37% | *INF*<br>36% | *ISO2*<br>25% | *RA*<br>x-bar = 36<br>±6.65SEM |
| 2<sup>nd</sup> Census<br>(Day 8) | *LP*<br>97% | *OLP*<br>91% | *DP*<br>76% | *MD*<br>70% | *ISO2*<br>67% | *S*<br>65% | *D*<br>64% | *ISO1*<br>60% | *INF*<br>13% | *PR*<br>1% | *RA*<br>x-bar = 159<br>±25.6SEM |

### Aphid polyphenism

The proportions of alates were significantly affected by aphid density in the microcosm (LRχ^2^_(1,48)_ = 6.49, P = 0.011); both the predator-associated risk treatment and the interaction between the treatment and aphid density had highly significant effects on alata production (LRχ2_(10,48)_ = 180.34, P < 0.0001) and (LRχ^2^_(10,48)_ = 42.32, P < 0.0001), respectively (see Supplementary Tables S4 for model details). Alate proportions differed across the predator-associated risk treatments. There were no alates to report under (*LP*), different from the unexpected percentage under (*OLP*) (x-bar = 2.8% ±2.78SEM).

More surprisingly, we found the highest morph proportion (x-bar = 16.7% ±5.3SEM) under (*ISO2*) (fed ladybirds left with no nourishment in the sachet), but (*ISO1*) (ladybirds enveloped with an aphid feed) resulted in a considerably small percentage of alates (x-bar = 0.7% ±0.7SEM), (Fig. 4). The proportion of alates was (x-bar = 1.5% ±0.4SEM) under the control (*RA*), attributable to crowding.

**Fig. 4.**
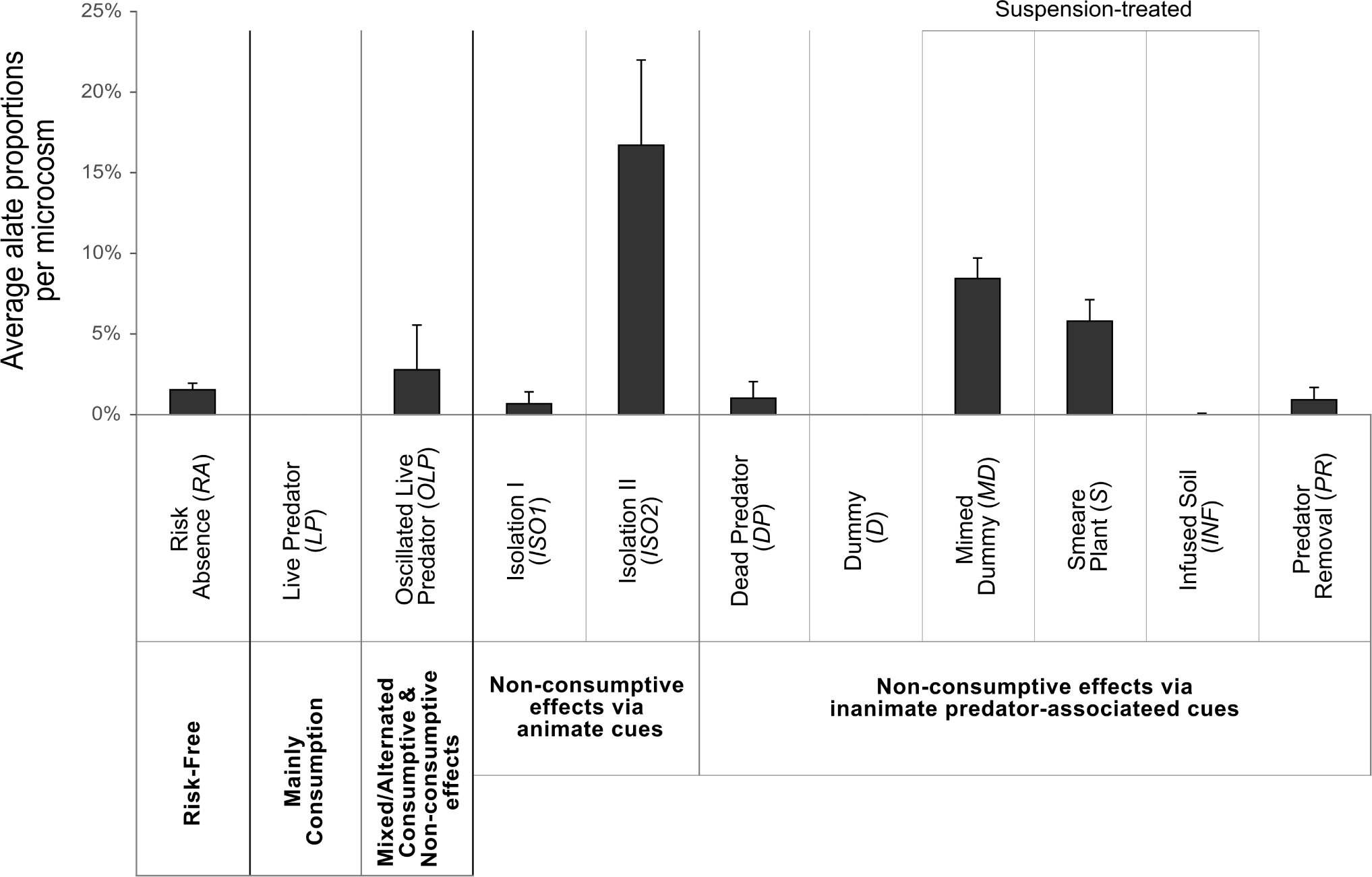
Aphid polyphenism under predator-associated risk treatments. Aphid polyphenism (distinct phenotypic plasticity of green peach aphid *M. persicae*) in response to the risk treatments associated with ladybird *C. septempunctata* is displayed as the mean percentage of alates (±SEM) relative to total aphid numbers in the microcosm on Day 8.

Amongst the inanimate risk-cue treatments, alates proportions were the highest in the presence of the *Mimed Dummies (MD*) (x-bar = 8.4% ±1.3SEM), followed by the *Smeared Plant* (*S*) (x-bar = 5.8% ±1.3SEM), both of the suspension-treated group. Conversely, with notable overall aphid reproductive success, the second smallest proportion of alates across all the risk treatments was recorded under (*INF)* (x-bar = 0.04% ±0.04SEM). Moreover, aphid reproductive success, on Day 8, was 75% lower under (*DP*) compared to (*PR*), yet counterintuitively, these non-consumptive risk treatments resulted in almost identical proportions of alates (x-bar = 1% ±1.02SEM) and (x-bar = 0.9% ±0.76SEM), respectively. Lastly, the risk imposed by the dummy (*D*) did not elicit any production of alates, (Fig. 4), (see also Supplementary *Note 1* including Tables S5-S7 for a complementary approach on analysing the effects of the different types of risk associated with the coccinellid predator).

## Discussion

Using a model agroecosystem of an important parthenogenetic plant pest raised on a significant crop, this work addresses repressive effects of fear of predation that have not been previously sufficiently investigated in aphid-predator systems. Intimidation by a range of different consumptive and/or non-consumptive effects associated with a coccinellid predator resulted in remarkably differential and time-limited impedance of the reproductive success of green peach aphid, with variable alata production (polyphenism); particularly, novel uses of predator corpses, cued plants, and cued predator dummies had strong impacts on the aphid population.

### Reproductive success and polyphenism under risk

Compared to the risk-free treatment (*RA*), *Live Predator* (*LP*) and *Oscillated Live Predator* (*OLP*) strongly impeded the reproductive success of *M. persicae*. The diminished aphid population in (*LP*) was the result of constant consumption mixed with possible non-consumptive effect (prey intimidation) (Preisser et al. 2005), whilst the predator foraged for food within the microcosm. The treatment (*OLP*) presented a complex periodic mixed-effect challenge as episodes of consumption accompanied by non-consumptive effects were interposed by predator time-outs (prey intimidation by predator-borne cues only) where the predator marks, bearing its semiochemicals, remained on the plants; different from (*LP*), this idiosyncrasy of variation in (*OLP*) induced more alates. It could be argued, therefore, that temporal variation and alternation of risk states (consumptive and non-consumptive) can induce various adaptive prey responses (Weisser, et al. 1999; Kunert and Weisser 2003; Keiser 2012; Keiser and Mondor 2013). Prey may reduce investment in and display of anti-predator behaviours under episodes of high or prolonged risk exposure (McNamara and Houston 1987; Lima and Bednekoff 1999; Hamilton and Heithaus 2001; Kotler et al. 2004), whilst the frequency of risk exposure may influence prey decisions (Lima and Bednekoff 1999) and shape prey propensities for escape and survival (Mackay and Wellington 1975; Lima and Bednekoff 1999; Keiser and Mondor 2013); the extent of the influence of risk on prey fitness and population dynamics has been proposed to be dependent on prey physiology as well as on temporal variability of risk (Trussell et al. 2011; Matassa and Trussell 2014). Our findings suggest that examining prey responses to pulsated variable intensities of predation risk (Lima and Bednekoff 1999; Trussell et al. 2011) should receive further investigation in future studies.

The non-consumptive effects of the animate risk associated with the isolated ladybird through its mere presence in the microcosm (*ISO2*) had a significant negative effect on the aphid population. Although the reproductive success was lower under (*ISO1*) than (*ISO2*) according to the 1^st^ census, the pattern of difference was surprisingly reversed in the 2^nd^ census, with more decrement of aphid numbers under (*ISO2*), with a remarkable ∼25-fold (i.e., 96%) increase in alata production. The aphid feed under (*ISO1*) was consumed alive in the micro-porous sachet poised in the vicinity of the target aphid clone; by Day 4, the ladybird having consumed the feed in isolation, there were seemingly plenty of aphid alarm signals diffusing from the sachet hence resulting in (*ISO1*) having a considerable impact. By contrast, up to the same day, the ladybird in (*ISO2*) was presumably emitting semiochemicals (Norin 2009) but that was not backed up enough by aphid alarm pheromone to result in a clear repressive effect. Afterwards, on Day 8, the ladybird in (*ISO2*) became increasingly hungry and might have been, therefore, emitting more intensive bursts of semiochemicals that triggered an alarm state in the clone and induced the production of alates; whereas, the effects of (*ISO1*) were noticeably weaker perhaps because there were no conspecifics left to be preyed upon by then and thus not enough a trigger of the alarm state. But, during the brief feeding period in (*ISO2*), it was also likely that the ladybird became smeared with cornicle secretions produced by stock aphids (the same clone as the experimental one); alarm signals might have emanated from the smeared secretions, after the ladybirds were re-introduced into the experimental units (Mondor and Roitberg 2004; Tamai and Choh 2019), and contributed to the induction of polyphenism and the population diminution via an extra non-consumptive effect. As such, the difference in aphid reproductive success between those animate risk treatments might be partly explained by the higher percentages of winged offspring, entailing reduced fecundity for the advantage of dispersal under (*ISO2*). More investment in dispersal, as an anti-predator defence, can be read as a transgenerational adaptive tactic (Keiser 2012; Keiser and Mondor 2013; Cabej 2019), but the inducible defence can be costly in the sense that it implicates a lower reproductive success due to a decreased production of fecund apterae (Mackay and Wellington 1975; Dixon 1998; Dixon and Agarwala 1999; Ingerslew and Finke 2017); survival of the clone may increase in the long term because of dispersion (Francke et al. 2008; Keiser 2012). Our findings imply that the prey-originated and the predator-originated cues might be seemingly intertwined in their collective effect on the aphid population, as the impact of the non-consumptive effects of the mere presence of the predator may need coupling with aphid alarm pheromones for the effect of the “scent of fear” (Leather 2015) to heighten (Ingerslew and Finke 2017); see also (Kats and Dill 1998; Fievet et al. 2009). Could it be, however, that the longer the isolated predator starves, up to a threshold, the more effective the release of the alert/alarm cues (eliciting flight defence) in the vicinity of the prey population?

The non-consumptive effects of the animate risk cues were by and large critical in impeding aphid reproductive success and generally of similar magnitudes to the non-the consumptive effects of the inanimate ones. Our results support earlier findings on the disruptive or suppressive effects of non-consumptive predation risk on aphid population dynamics (Nelson and Rosenheim 2006; Nelson 2007) including the effects of dead conspecifics (Fievet et al. 2008), predator bio-signature (Steffan and Snyder 2010; Ninkovic et al. 2013), and impaired predators (Nelson et al. 2004; Nelson and Rosenheim 2006). The population-diminishing effects of the inanimate risk cues pertaining to (*DP*), (*D*), (*MD*), and (*S*) were largely persistent by the 2^nd^ census, suggesting an extended negative impact on aphid reproductive success against time. In stark contrast, the effects of the soil-infused (*INF*) and the predator removal (*PR*) were short-lived and weakened after Day 4. It should be highlighted that by the 2^nd^ census, the effect of the predator replicas in the *Dummy* treatment (*D*), representing predator visual mimicry, became augmented in impeding aphid reproductive success after the daubing of the dummy with the predator suspension bearing olfactory cues in the *Mimed Dummy* treatment (*MD*). Yet the effect of (*MD*) could not surpass that of the predator corpse in (*DP*) across censuses. Eventually, (*DP*) had largely a stronger negative influence on the reproductive success than the predator-mimicking treatments (*Dummy, Mimed Dummy, Smeared Plant*, and *Soil Infused*) and the animate-risk treatments (*ISO1*) and (*ISO2*). This possibly pertains to more reliable visual and olfactory plus tactile cues the predator corpses in (*DP*), followed by the mimed dummies in (*MD*), bore in contact with the target aphids. The rest of the non-consumptive risk treatments, which mostly entailed either visual or olfactory signals, were not as effective as the predator corpses and the dummies. This receives credence from the findings documented by Ingerslew and Finke (2017) on the palpation of aphids by parasitoid wasps, suggesting an under-studied non-consumptive negative tactile effect on aphid fitness that might be applicable for other aphidophagous natural enemies. The combination of several aphid perceptions (Ben-Ari and Inbar 2014) of distinct concurrent risk cues could be the decisive factor behind the superlative repressive effect of (*DP*) in the long run. Overall, our findings comparatively showcase that dead predators and *biomimetic* synthetics along with manipulating the non-consumptive effects of isolated predators may, in novel fashions, lead to tangible pest control results.

Not only can polyphenism be induced by aphid density (Dixon 1998), but also by a variety of other factors spanning residing in exhausted resources and the presence of natural enemies (Dixon 1998; Kunert et al. 2005; Creel and Christianson 2008); previous studies showed that ladybird traces might, as well, trigger the production of alates (Dixon and Agarwala 1999). Our results indicate that intimidation signals by the animate risk in (*ISO2*) and the inanimate risks in (*M*) and (*S*) might have played a critical role in the induction of the winged morphs as a phenotypically plastic response the risk, irrespective of the smaller corresponding populations suffering from repression pressure of the ranks 4^th^, 5^th^, and 6^th^, respectively, as per the 2^nd^ census. However, Purandare et al. (2014) demonstrated in a study on transgenerational polyphenism in pea aphid that the tracks of a coccinellid predator induced the least number of dispersive morphs if compared with the control or the effect of crowding; the cited authors suggested that these polyphenic responses might be dependent on the intensity and sufficiency of predation risk cues. Nevertheless, the proportions of winged offspring continued to differ under the influence of non-consumptive effects in our study. For example, the treatments (*MD*) and (*S*) led to a greater number of alates while impeding reproductive success. As the dispersive winged phenotype is less fecund than the wingless (Roitberg et al. 1979, Dixon 1998, Kunert and Weisser 2003), alata production can also account for the reduced aphid densities under risk in (*MD*) and (*S*) due to increased investment in the generation of alates. In comparison, under (*INF*) and (*PR*), the aphid population thrived (yet produced negligible to relatively small numbers of alates). This implies that alarm signals (pheromones) and risk cues (semiochemicals), not crowdedness in these cases, were decisive in inducing polyphenism as a defence to flee the risk.

In other words, our results suggest that the mimics (dummies, mimed dummies, and the cued plant) might have ‘tricked’ the aphids as the decoys seemed to have been perceived as a real threat that altered the transgenerational phenotypic plasticities and and adaptations (Keiser 2012) of the exposed aphids. Concomitant modified reproduction plus alata production (Nelson et al. 2004, Nelson 2007; Khudr et al. 2017) seem to be contextual anti-predator tactics that may change relevant to aphid physiological state and the state and the nature of risk (McAllister et al. 1990; Mousseau and Fox 1998; Weisser et al. 1999; Villagra et al. 2002; Hu et al. 2018), the type and frequency of risk exposure (Sih et al. 2000; Matassa and Trussell 2014). Again, the anti-predator responses involve energetic and physiological tolls (Creel and Christianson 2008) on the reproduction of the clonal prey population (Dixon and Agarwala 1999; Nelson et al. 2004; Mondor et al. 2005; Nelson 2007) wherever aphids encounter predators and predation risk (Nelson et al. 2004; Nelson and Rosenheim 2006; Nelson 2007). Still, other possible explanations for our results might be partly assignable to mortality as a result of predator-induced stress, as suggested by McCauley et al. (2011), and/or reproductive costs following abrupt changes in diet and disruption of feeding time (Nelson 2007). All in all, the induction of polyphenism by the treatment including mimetic dummies or treated host plants with predator-borne cues unveiled extreme phenotypic plasticity (Mondor et al. 2005; Whitman and Agrawal 2009) in green peach aphid as means of escaping novel threats (Grostal and Dicke 1999). We can assume, therefore, that the influence of fear of predation on aphid fitness might be pronounced in environments with frequent disturbances and disproportionate intensities of localised environmental risk. This is concordant with the findings reported by Lin and Pennings (2018) suggesting that aphid control by ladybirds can be strong on fine spatial scales, as in our study system, where the aphid-ladybird interactions have been shown to be more prominent (Lin and Pennings 2018).

Based on the risk allocation hypothesis (Lima and Bednekoff 1999), we propose that the variation in responses to the cocktail of non-consumptive effects in our study can be largely attributed to *M. persicae*’s ability to perceive, identify and respond to different kinds, intensities and frequencies of environmental risks by relying on combined modalities (Ben-Ari and Inbar 2014). As they assesscues from their embedding contexts, aphids adjust their progenies phenotypes to adapt to change and procreate (Dixon 1998; Dixon and Agarwala 1999; Keiser 2012; Ben-Ari and Inbar 2014); less accurate or wrong assessment of risk can lead to subsequent maladaptive or delayed decisions ending with fitness loss (Mackay and Wellington 1975; Nelson 2007; Keiser 2012), but see (Tamai and Choh 2019). But, it can be argued that the non-consumptive risk treatments in our work presented variably unreliable cues that led to perceiving the embedding contexts by the aphids of the female-only clone as rather inhospitable; this would have created, in turn, a varying state of sensory ambiguity that altered prey’s decisions and consequently hampered aphid fitness (Koops 2004). For instance, the notable difference between the impacts of (*PR*) and (*S*) on the aphid population lies in the fact that the intensity of the predator-borne cues smeared on the plant in (*S*) was up to or above a threshold necessary to elicit polyphenism as plastic anti-predator defence causing aphid reproductive success to decelerate. The impact of (*S*) was accentuated anthropogenically by daubing (smearing) unlike the (*PR*) treatment that just imitated a natural process when a predator marks a plant whilst searching for prey. Further, the short-lived effects of (*INF*) and (*PR*) conform to the work provided by Koops (2004), suggesting that the acquisition of ambiguous information may have a shorter temporal influence on prey decision making than the information acquired from reliable cues. Notwithstanding, the weak effects of the (*INF*) and (*PR*) in the long run could also be on account of risk habituation; it has been shown that prey responsiveness to predation risk may decrease due to habituation after continuous exposure to non-lethal cues (Shalter 1984; Holomuzki and Hatchett 1994).

At any rate, we did not observe any clear direct avoidance of the cue-treated plants in the microcosm by aphids, but the phenotypic plasticity, denoting dispersal, was detectable as described above (see Supplementary *Note 2*). Further, *M. persicae* also varied its inducible defences under (*ISO2*) of prey intimidation by animate risk, as there was less investment in the production of apterae, increased maternal conditioning of offspring into alates, and relatively more aphids abandoned and/or dropped off their host plants (see also Supplementary *Note 3* including Table S8 and Fig. S1 for a spotlight on a complementary off-plant behaviour).

In summary, amid research scarcity on the effects of indirect and imitative stimuli associated with aphidophagy, our assay provides empirical evidence of the negative impact of non-consumptive effects associated with a coccinellid predator on a clonal population of an important aphid species, stressing the importance of widening the scope of investigation of prey intimidation in predator-prey systems. Interestingly, the isolated starved predator led to unparalleled intensified induction of phenotypic plasticity, and the effects of the inanimate cues resulted in better or similar negative impacts on aphid reproductive success when compared to those of the animate risk (i.e., the presence of an isolated predator). We also found that certain risk effects may have a variable longevity in a context-dependent fashion; specific types of novel prey intimidation by dead predators and mimed dummies were persistent in effect and considerably more efficient than others (e.g., soil infusion with predator-borne cues). Our findings have the potential to provide unorthodox eco-friendly means to protect agroecosystems from pest irruptions *in situ* and *in natura*.

## Declarations

## Acknowledgements

Special thanks go to Dr Inga Mewis (JKI, Berlin) for providing an initial sample of the *M. persicae* clone used in this study, and to Dr Reinmar Hager for hosting MSK while finalising the manuscript. Many thanks to Hayley Craig for the insights and comments on improving the manuscript. This work received funding from the Freie Universität Berlin for MSK. OYB was funded by the DRS Point Fellowship, Freie Universität Berlin.

