## Supplementary for "Live and inanimate predator-associated cues suppress the population of sap-feeding prey and induce polyphenism"

**Running title:** Predator-associated risk cues suppress prey population

**Table S1. Experimental design.** The table explains the structure and concept of the experimental design with brief description of the charismatics of each of the risk treatments associated with the aphidophagous predator ladybird ladybird *Coccinella septempunctata*; the target prey species was a clone of green peach aphid *Myzus persicae* (see Fig. 1, main text for a diagram of the design).

| Risk Type | Risk Treatment | Treatment details |
|---|---|---|
| Risk-free | Risk Absence (*RA*) | Aphids were alone in the microcosm. |
| Consumptive mixed with non-consumptive effect | Live Predator (*LP*) | Ladybirds were individually introduced per microcosm and remained enclosed for the full treatment time. The risk effect here was consumptive accompanied by non-consumptive effect resulting from foraging. |
| Consumption, mixed and alternated with non-consumptive effect | Oscillated Live Predator (*OLP*) | The predator was left in the enclosure (microcosm) for 2 days followed by one day 'out of microcosm' and so forth periodically until the end of the experiment. The risk effect here was consumptive accompanied by non-consumptive effect resulting from foraging (ladybird time-ins), and non-consumptive (ladybird time-outs, predator marks, footprints, excretions). |
| Non-consumptive Animate Cues | Isolation I (*ISO1*) | Each ladybird was enveloped with 20 aphids to feed on, in a fine-mesh sachet of about 3x3cm in diameter attached to one plant per microcosm. Ladybird individuals were left enveloped continuously throughout the treatment. The risk effect here was non-consumptive via isolated non-starved predator presence and emanating alarm pheromones from the consumed aphid conspecifics confined in the predator sachet. |
| | Isolation II (*ISO2*) | Ladybirds were individually enveloped, in the above mentioned attachable sachets of Isolation-I, but without aphids. On day 4, during aphid census, ladybirds were captured briefly and fed with aphids from the culture in separated plastic containers before they were re-enveloped in the corresponding sachets. The risk effect here was non-consumptive and constant, starved predator mere presence. |
| Non-consumptive Inanimate Cues | Dead Predator (*DP*) | Fourteen ladybirds were frozen. Then, two randomly selected ones were transferred into each pot; one ladybird was placed on the ground close to the stem, while the other was carefully tied to the stem ~3cm above soil surface, using a sewing kit. The risk effect here was non-consumptive bearing visual, tactile and olfactory cues. |
| | *Dummy* (*D*) | Waterproofed mock ladybirds, mimicking ladybird, were used. The dummies were placed on different vertical levels (as in the treatment *Dead Predator*). The risk effect here was non-consumptive (visual mimicry). |
| | *Mimed Dummy* (MD) Suspension-treated | Dummies were bound to one plant in the enclosure as described in the *Dummy* treatment above and carefully smudged several times with the dead-ladybird suspension. The risk effect here was non-consumptive (visual and olfactory/chemical mimicry). |
| | *Smeared Plant* (*S*) Suspension-treated | The leaves and stem of one plant in each corresponding microcosm were generously daubed with the dead-ladybird suspension. The risk effect here was non-consumptive (olfactory/chemical cues). |
| | *Soil Infused* (*INF*) Suspension-treated | The dead-ladybird suspension was infused into the soil at varying depths adjacent to the plant root system. This created a fine surrounding 'prey intimidation zone' in and on the soil surrounding the root system and the stem base of the treated plant. The risk effect here was non-consumptive (mainly indirect olfactory/chemical cues). |
| | *Predator Removal* (*PR*) | For 48 h, one ladybird adult was left with one plant only per microcosm in order to have the plants marked by predator bio-signature. Subsequently, after ladybird removal, a fresh plant (untreated) was carefully transferred to each microcosm with a mass of soil surrounding the roots. The risk effect here was non-consumptive (semiochemicals, predator tracks, traces and execrations). |

**Table S2. Summary of the generalised linear mixed-effect model (GLMM) on aphid reproductive success under predator-associated risk treatment.** Details are provided regarding the GLMM (specified in the main text) investigating aphid reproductive success of a clone of green peach aphid *M. persicae* in response to the 11-level risk treatment associated with ladybird *C. septempunctata*. The risk treatments were: Live Predator (*LP*), Oscillated Live Predator (*OLP*), Isolation I (*ISO1*), Isolation II (*ISO2*), Dead Predator (*DP*), Dummy (*D*), Mimed Dummy (*MD*), Smeared Plant (*S*), Soil Infused (*INF*), and Predator Removal (*PR*); Risk Absence (*RA*) was the model reference. See the main text methods for model specifications. Significant results are shown in bold.

| Fixed effects Risk Treatment | Estimate | Std. Error | t value | P |
|---|---|---|---|---|
| *LP* | 0.157127 | 0.031299 | 5.020 | **<0.0001** |
| *OLP* | 0.123406 | 0.023361 | 5.282 | **<0.0001** |
| *ISO1* | 0.023161 | 0.009910 | 2.337 | **0.019** |
| *ISO2* | 0.024406 | 0.009557 | 2.554 | **0.011** |
| *DP* | 0.041878 | 0.011459 | 3.655 | **0.0003** |
| *D* | 0.025288 | 0.010172 | 2.486 | **0.013** |
| *MD* | 0.033753 | 0.011282 | 2.992 | **0.003** |
| *S* | 0.028683 | 0.010096 | 2.841 | **0.004** |
| *INF* | 0.008411 | 0.008745 | 0.962 | 0.336 |
| *PR* | 0.010501 | 0.009674 | 1.086 | 0.278 |

**Table S3. Aphid reproductive success, multiple post-hoc comparisons under predator-associated risk treatment.** The outcome is provided regarding the multiple post-hoc pairwise comparisons (Tukey's HSD) following the GLMM testing the reproductive success of green peach aphid *M. persicae* in response in response to the 11-level risk treatment associated with ladybird predator *C. septempunctata*. The risk treatments were: Live Predator (*LP*), Oscillated Live Predator (*OLP*), Isolation I (*ISO1*), Isolation II (*ISO2*), Dead Predator (*DP*), Dummy (*D*), Mimed Dummy (*MD*), Smeared Plant (*S*), Soil Infused (*INF*), and Predator Removal (*PR*); Risk Absence (*RA*) was the model reference. Only significant results (bold) are displayed, confidence level used: 0.95.

| Predator-associated risk treatment comparative pairs | estimate | SE | z.ratio | P |
|---|---|---|---|---|
| *RA versus LP* | -0.157126841 | 0.031298771 | -5.020 | **< 0.0001** |
| *RA versus OLP* | -0.123405813 | 0.023361270 | -5.282 | **< 0.0001** |
| *RA versus DP* | -0.041877753 | 0.011458563 | 3.655 | **0.012** |
| *LP versus ISO1* | 0.133966126 | 0.031727700 | 4.222 | **0.001** |
| *LP versus ISO2* | 0.132720633 | 0.031649290 | 4.193 | **0.001** |
| *LP versus DP* | 0.115249088 | 0.032232980 | 3.576 | **0.015** |
| *LP versus D* | 0.131838923 | 0.031804014 | 4.145 | **0.002** |
| *LP versus MD* | 0.123373409 | 0.032142375 | 3.838 | **0.006** |
| *LP versus S* | 0.128443476 | 0.031777424 | 4.042 | **0.003** |
| *LP versus INF* | 0.148715973 | 0.031359372 | 4.742 | **0.0001** |
| *LP versus PR* | 0.146625419 | 0.031560154 | 4.646 | **0.0002** |
| *OLP versus ISO1* | 0.100245098 | 0.023875871 | 4.199 | **0.001** |
| *OLP versus ISO2* | 0.098999605 | 0.023799138 | 4.160 | **0.002** |
| *OLP versus DP* | 0.081528060 | 0.024545767 | 3.321 | **0.036** |
| *OLP versus D* | 0.098117895 | 0.023974213 | 4.093 | **0.002** |
| *OLP versus MD* | 0.089652381 | 0.024400605 | 3.674 | **0.011** |
| *OLP versus S* | 0.094722448 | 0.023935110 | 3.957 | **0.004** |
| *OLP versus INF* | 0.114994945 | 0.023374950 | 4.920 | **< 0.0001** |
| *OLP versus PR* | 0.112904391 | 0.023598335 | 4.784 | **0.0001** |

**Table S4. Details of the generalised linear model (GLM) on aphid polyphenism under predator associated risk treatments.** Details are provided regarding the GLM investigating alata production of a clone of green peach aphid *M. persicae* in response to the 11-level risk treatment associated with ladybird *C. septempunctata*, and numerical pressure of the total aphid numbers (density) in the microcosm that was used as a covariate. The risk treatments were: Live Predator (*LP*), Oscillated Live Predator (*OLP*), Isolation I (*ISO1*), Isolation II (*ISO2*), Dead Predator (*DP*), Dummy (*D*), Mimed Dummy (*MD*), Smeared Plant (*S*), Soil Infused (*INF*), and Predator Removal (*PR*); Risk Absence (*RA*) was the model reference. Significant results are displayed in bold.

| Analysis of Deviance Table (Type II tests) | LR$\chi^2$ | Df | Pr(>Chisq) |
|---|---|---|---|
| Aphid density | 6.49 | 1 | **0.011** |
| Risk Treatment | 180.34 | 10 | **<0.0001** |
| Risk Treatment × Aphid density | 42.32 | 10 | **<0.0001** |

(Dispersion parameter for quasipoisson family taken to be 2.150198)
Null deviance: 616.80 on 69 degrees of freedom
Residual deviance: 103.65 on 48 degrees of freedom

***Note 1:*** *Complimentary analysis of aphid reproductive success and polyphenism under different types of predator-associated risks*

Here we show the analysis of aphid reproductive success and polyphenism from the standpoint of these traits being altered in response to the different predator-associated risk types designated into comparable categories (see Supplementary Tables S5-S7).

A generalised linear mixed effect model (GLMM) was used in R (R Core Team 2016) to examine aphid reproductive success (total aphid in the microcosm), function 'glmer', 'bobyqa' optimisation; gamma family (due to the non-normal shape of distribution, confirmed by Shapiro test, and high positive skewness of the data histogram), using R package 'lme4' (Bates et al. 2015). There were two counts (on Day 4, 1$^{st}$ census) and (on Day 8, 2$^{nd}$ census). The microcosm (plants, aphids and their pots enclosed with mesh as described in the main text) was nested within the count days and randomised in the model; the main effects of the models were respectively revealed using an Anova command, R packages 'car' (Fox and Weisberg 2011). The predator-associated risks (fixed effects) corresponding to the 11-level treatment were as follows: Absence of risk of the treatment (*RA*) was the risk-free control; consumption accompanied by non-consumptive risks under the treatment Live Predator (*LP*), consumptive risk mixed/alternated with non-consumptive risk under the treatment Oscillated Live Predator (*OLP),* non-consumptive risk via using animate cues under the treatments of an isolated predator with aphid feed [Isolation I (*ISO1*)] and an isolated predator without the feed [Isolation I (*ISO2*)], and non-consumptive risk via using inanimate cues under the treatments of Dead Predator (*DP),* predator replica [Dummy (*D*)], Mimed Dummy (*MD*), [Smeared Plant (*S*)], [Soil Infused (*INF*)], and [Predator Removal (*PR*)]. The dummies in (*MD*) and the cued-plant in (*S*) where treated/prompted with predator suspension (specified in the main text) bearing ladybird-borne cues; in the treatment (*INF*) the suspension was injected into the soil around one plant in the microcosm; in the treatment (*PR*) only one plant had a previous interaction with the predator and hence was marked by its bio-signature (e.g., track, excretions, and semiochemicals) before being transferred into the experimental microcosm. We also applied a posthoc pair-wise multiple comparison test (Tukey's HSD), using R package 'lsmeans' (Lenth 2016).

As shown in (Fig. 3 of the main text), aphids suffered a clear loss in reproductive success that was contextual and contingent upon the type of the imposed predator-associated risk. The latter highly influenced aphid reproductive success in the microcosm ($\chi^2_{(4,268)} = 50.90$, $P < 0.0001$), where, according to the GLMM summary, all risk types had differential significant effects as follows: mainly consumptive risk via the treatment Live Predator (*LP*) (P = 0.023), mixed/alternated consumptive and non-consumptive risks via Oscillated Live Predator (*OLP*) (P < 0.0001), non-consumptive risk via animate cues of the treatment Isolation I (*ISO1*) and Isolation II (*ISO2*) (P = 0.008), non-consumptive risk via inanimate cues of the treatments Dead Predator (*DP*), *D* (*D*), Mimed Dummy (*MD*), Smeared Plant (*S*), Soil Infused (*INF*), and Predator Removal (*PR*) (P = 0.002), (see Supplementary Table S5 for details of the GLMM model, and Table S6 for posthoc multiple pairwise comparisons).

We also investigated the effects of the aforementioned predator-associated risk types on the induction of alates as the winged morphs were counted on Day 8 at the end of the experiment. We applied a generalised linear model (GLM) with a quasipoisson family (due to over-dispersion and non-normality [confirmed by Shapiro test]), using the R package 'multcomp' (Hothorn et al. 2008). The effects were: 1) The 11-level risk treatment, where "Risk Absence (*RA*)" was the model baseline, 2) Aphid total numbers per microcosm (aphid density), and 3) the interaction between these effects. The main effects of the models were revealed using an Anova command, R packages 'car' (Fox and Weisberg 2011).

The GLM showed that the type of predator-associated risk highly influenced alata production ($LR\chi^2_{(4,60)} = 27.07$, $P<0.0001$), so did aphid density ($LR\chi^2_{(1,60)}= 8.63$, $P = 0.003$); but the interaction between these two predictors was marginally significant ($LR\chi^2_{(4,60)} = 9.5$, $P = 0.0497$) (see main text Fig. 4, and Supplementary Table S7 for details of the GLM model).

**Table S5. Summary of the generalised linear mixed-effect model (GLMM) on aphid reproductive success under different predator-associated risk types.** Details are provided regarding the GLMM investigating aphid reproductive success (total numbers of aphids in the microcosm) of a clone of green peach aphid *M. persicae* in response to different risk types associated with ladybird *C. septempunctata*. The risk types were: Absence of risk of the treatment (*RA*) as the risk-free control; consumption accompanied by non-consumptive effect was the risk under the treatment Live Predator (*LP*), consumptive risk mixed/alternated with non-consumptive risk under the treatment Oscillated Live Predator (*OLP),* non-consumptive risks due to using animate cues under the treatment of an isolated predator with an aphid feed [Isolation I (*ISO1*)] and of an isolated predator without an aphid feed [Isolation I (*ISO2*)], and non-consumptive risks using inanimate cues under Dead Predator (*DP*), predator replica [Dummy (*D*)], Mimed Dummy (*MD*), Smeared Plant (*S*), Soil Infused (*INF*), and Predator Removal (*PR*). Significant results are shown in bold. See the main text Methods for model specifications.

| Fixed effects<br>Risk Types | Estimate | Std. Error | t value | P |
|---|---|---|---|---|
| Consumption accompanied by non-consumptive risks of (*LP*) | 0.158725 | 0.031135 | 5.098 | **<0.0001** |
| Consumptive risk, accompanied by and alternated with non-consumptive risk of (*OLP*) | 0.126018 | 0.023551 | 5.351 | **<0.0001** |
| Non-consumptive risk via using animate cues | 0.026665 | 0.010039 | 2.656 | **0.008** |
| Non-consumptive risk via using inanimate cues | 0.027454 | 0.009039 | 3.037 | **0.002** |

| AIC | BIC | logLik | deviance | df.resid |
|---|---|---|---|---|
| 2095.1 | 2124.1 | -1039.6 | 2079.1 | 268 |

**Table S6. Aphid reproductive success, multiple post-hoc comparisons under different predator-associated risks.** Details are provided regarding the multiple post-hoc pairwise comparisons (Tukey's HSD) following the GLM testing aphid reproductive success of a clone of green peach aphid *M. persicae* in response to different risk types associated with ladybird *C. septempunctata*. The risk types were: Absence of risk of the treatment (*RA*) as the risk-free control; consumption accompanied by non-consumptive effect was the risk under the treatment Live Predator (*LP*), consumptive risk mixed/alternated with non-consumptive risk under the treatment Oscillated Live Predator (*OLP),* non-consumptive risks due to using animate cues under the treatment of an isolated predator with an aphid feed [Isolation I (*ISO1*)] and of an isolated predator without an aphid feed [Isolation I (*ISO2*)], and non-consumptive risks using inanimate cues under Dead Predator (*DP*), predator replica [Dummy (*D*)], Mimed Dummy (*MD*), Smeared Plant (*S*), Soil Infused (*INF*), and Predator Removal (*PR*). Significant results are shown in bold. See the main text Methods for model specifications. Only significant (bold) or marginally significant results are displayed, confidence level used: 0.95.

| Contrast | Estimate | SE | z.ratio | P |
|---|---|---|---|---|
| Risk-free of (*RA*) *versus* Consumption accompanied by non-consumptive risks | -0.1587 | 0.0311 | -5.098 | **<0.0001** |
| Risk-free of (*RA*) *versus* Consumptive risk, accompanied by and alternated with non-consumptive risk | -0.1260 | 0.0236 | -5.351 | **<0.0001** |
| Risk-free of (*RA*) *versus* Non-consumptive risk via using animate cues | -0.0267 | 0.0100 | -2.656 | 0.061 |
| Risk-free of (*RA*) *versus* Non-consumptive risk via using inanimate cues | -0.0275 | 0.0090 | -3.037 | **0.020** |
| Consumption accompanied by non-consumptive risks *versus* Non-consumptive risk via using animate cues | 0.1321 | 0.0308 | 4.281 | **0.0002** |
| Consumptive risk, accompanied by and alternated with non-consumptive risk *versus* Non-consumptive risk via using inanimate cues | 0.1313 | 0.0305 | 4.305 | **0.0002** |
| Consumptive risk, accompanied by and alternated with non-consumptive risk *versus* Non-consumptive risk via using animate cues | 0.0994 | 0.0231 | 4.296 | **0.0002** |
| Consumptive risk, accompanied by and alternated with non-consumptive risk *versus* Non-consumptive risk via using inanimate cues | 0.0986 | 0.0226 | 4.362 | **0.0001** |

| AIC | BIC | logLik | deviance | df.resid |
|---|---|---|---|---|
| 2094.3 | 2145.0 | -1033.2 | 2066.3 | 262 |

**Table S7. Details of the generalised linear model (GLM) on aphid polyphenism under different predator-associated risks.** Details are provided regarding the details of the GLM investigating alata production of a clone of green peach aphid *M. persicae* in response to different risk types associated with ladybird *C. septempunctata*, and numerical pressure of the total aphid numbers in the microcosm (density) that was used as a covariate. Risk Absence (*RA*) was the control. Risk Absence (*RA*) was the control. Consumption accompanied by non-consumptive effect was under (*LP* [Live Predator]), consumption, accompanied by and alternated with non-consumptive effect, was under (*OLP* [Oscillated Live Predator]), non-consumptive effects using animate cues were under (*ISO1* [Isolation I] and *ISO2* [Isolation II]), and non-consumptive effects using inanimate cues were under (*DP* [Dead Predator], *D* [Dummy], *MD* [Mimed Dummy], *S* [Smeared Plant], *INF* [Soil Infused], *PR* [Predator Removal]). See the main text methods for model specifications.

| Analysis of Deviance Table (Type II tests) | LR$\chi^2$ | Df | Pr(>Chisq) |
|---|---|---|---|
| Aphid density | 8.63 | 1 | **0.003** |
| Risk Type | 27.07 | 4 | **<0.0001** |
| Risk Type × Aphid density | 9.5 | 4 | **0.0497** |

(Dispersion parameter for quasipoisson family taken to be 5.693547)
Null deviance: 616.80 on 69 degrees of freedom
Residual deviance: 374.18 on 60 degrees of freedom

***Note 2:*** *The absence of avoidance behaviour*

From the main text Results (Fig. 3, Table 1, and Fig. 4), it was clear that aphid reproductive success plummeted and the production of alates (anti-predator defence) was relatively considerable under the risk treatments Smeared Plant (*S*) and Mimed Dummy (*MD*), but the reproductive success increased sharply under the treatments Soil Infused (*INF*) and Predator Removal (*PR*), both with poor production of alates irrespective of the higher aphid densities in the microcosm. We can easily glean from these findings that the predator-associated risk effects accompanied by aphid alarm pheromonal communication might have gradually weakened for specific risk stimuli hence the increased reproductive success and decreased polyphenism over time in the mentioned examples. However, we note that it is eye-catchy that across the treatments, we did not observe significant anti-predator avoidance behaviour of the plants treated with non-consumptive predator-associated stimuli (i.e., cued plants); this includes the cued plants in the inanimate risk treatments Dead Predator (*DP*), Dummy (*D*), Mimed Dummy (*MD*), Smeared Plant (*S*), and Soil Infused (*INF*). It can be argued that the generally diminished avoidance behaviour was considerably owing to the non-lethality of the risk cues used, which differentially impacted on aphid reproductive success and alata production, but were not certain enough to alter the behaviour of aphids (Koops 2004; Ferrari et al. 2010; Ben-Ari & Inbar 2014). Also, it might be argued that *Myzus persicae* might have got accustomed to those types of non-consumptive cues applied in our experiment (Holomuzki & Hatchett 1994; de Vos et al. 2010). In general, the negligible avoidance behaviour in our study resonates with the findings reported on by Costamagna et al. (2013) on the absence of avoidance behaviour in the soybean aphid *Aphid glycines.*

At any rate, *M. persicae*'s general propensity to aggregate through the applied risk with no discrimination against risk hotspots imposed costs on its population growth across certain risk treatments (see Fig. 3 of the main text); such costs might have emerged from prey intimidation (fear of predation) and as a consequence of induced stress (McCauley et al. 2011; LaManna & Martin 2016; Khudr et al. 2017) or predator semiochemicals (Norin 2009); the effect of these stressors could be escalated by aphid alarm pheromone (Keiser & Mondor 2013; Ingerslew & Finke 2016). Aphid responses as observed in our work are considered to be contextual in accord with the better or poorer aphid performance. However, have the prey fallen into the traps of not recognising an imminent danger due to risk cue ambiguity (Koops 2004; Ben-Ari & Inbar 2014), or made an error in the risk assessment and following colonisation decisions of a risky micro-habitat, the impact on their fitness and survival can be significant (Ben-Ari & Inbar 2014). This is expected, as well, to vary depending on prey life stages and specific adaptive response to predation risk (Dixon & Agarwala 1999; Keiser 2012; Keiser & Mondor 2013; LaManna & Martin 2016); (see also Supplementary *Note 3*).

***Note 3:*** *Off-plant distribution*

We examined, in R, the effects of the risk treatment associated with ladybird predator *C. septempunctata* on the propensity of green peach aphid *M. persicae* to drop-off their host plants as the proportions of off-plant aphids were calculated on Day 4 (1at census) and Day 8 ($2^{nd}$ census). There were 11 risk treatments as follows: Live Predator (*LP*), Oscillated Live Predator (*OLP*), Isolation I (*ISO1*), Isolation II (*ISO2*), Dead Predator (*DP*), Dummy (*D*), Mimed Dummy (*MD*), Smeared Plant (*S*), Soil Infused (*INF*), and Predator Removal (*PR*), (see main text Methods for further specifications). We applied a generalised linear model (GLM) with a quasipoisson family (due to over-dispersion and non-normality [confirmed by Shapiro test]), using the R package 'multcomp'. The effects were: 1) The 11-level risk treatment, where "Risk Absence (*RA*)" was the model baseline; the treatment was nested in the count day, 2) Aphid total numbers per microcosm (aphid density) as a covariate, and 3) the interaction between these effects. The main effects of the models were revealed using an Anova command, R packages 'car' (Fox and Weisberg 2011).

The GLM showed that the only the risk treatment and the interaction (aphid density x risk treatment x count day) were significant ($LR\chi^2_{(10,109)} = 32.62$, $P = 0.0003$) and ($LR\chi^2_{(11,109)} = 30.65$, $P = 0.003$), respectively (see Supplementary Table S8 for details of the GLM model, and Fig. S1). We observed negligible to small aphid numbers off plants on Day 4 and Day 8 under the effect of the treatment of 11 levels of predator-associated risk. Off-plant aphid proportions, in the case of Risk Absence (*RA*), were barely surprising due to a density effect (crowding), as the aphid population thrived in the optimal risk-free environment. A similar explanation can be provided for the case of Predator Removal (*PR*). Noticeably, when the live predator was free to forage in the microcosm, aphids did not wander off their host plants or were hunted by the ladybird if they did so. However, the escalated propensity to be off plant under the case of consumption-non-consumption alternation in (*OLP*) as well as in the case of the isolated starved predator of (*ISO2*) was apparent. This could be attributed to changes in the info-chemical alert and alarm in the clone due to the idiosyncrasies of the cues in the said treatments; with corresponding low aphid reproductive success, it is clear that crowdedness was not a factor therein but fluctuated intensities of risk was (see main text Methods and Results for further details). By contrast, it can be argued that although the risk cues under (*PR*) and (*INF*) were deficient in strength to repress the aphid population when compared to other risk tenements, the set-up of the said inanimate-risk-cue treatments might have altered the micro-climatic conditions of the aphid environment in way that led, in interaction with aphid density (crowding effect), to altered aphid behaviour and thus an induced plant abandonment in the long run (Dixon & Agarwala 1999; Nelson et al. 2004; Nelson 2007; Keiser & Mondor 2013).

Nevertheless, what attracts attention is that the propensity to go off plant was noticeable in the cases of the suspension-treated treatments (*S and MD*); olfactory cues in the former and visual plus to olfactory ones in the latter did hamper aphid reproduction and it appears that the risk cues were strong enough there to induce a fraction of the aphid population to abandon their host plants over time (Supplementary Table S8, and Fig. S1; see also Fig. 1 and Fig. 3 of the main text). Given the sedentary nature of *M. persicae* and a suggestion by Ingerslew & Finke (2016) that, unlike pea aphid *Acyrthosiphon pisum*, dropping off plant is not a favourable defence by *M. persicae* against predation risk, we advocate that when apterous aphids in our study did show dispersion by walking off plant it was tactical as well as contextual because usually dropping off-plant is a costly behaviour due to mortality risks being exposed to natural enemies *and also* a concurrent disruption of aphid necessary constant feeding (Nelson et al. 2004; Nelson 2007; Hoki et al. 2014). Our findings corroborate the norm that dispersion by alates, rather than off-plant dropping by apterae, would be the favoured anti-predator defence by *M. persicae* under predation risk. Still, the measured trait herein of abandoning the host plants should be guardedly used as a proxy for aphid escape behaviour (inducible anti-predator defence) when surveying the responses of this aphid species.

**Table S8. Aphid percentage off plant under predator-associated risk treatments.** A generalised linear model (GLM), as described above, was applied to test proportions of green peach aphid *M. persicae* that were found off plants upon two censuses [Day 4 and Day 8]) in response to the risk treatments associated with ladybird *C. septempunctata* comprising the following 11 levels: Live Predator (*LP*), Oscillated Live Predator (*OLP*), Isolation I (*ISO1*), Isolation II (*ISO2*), Dead Predator (*DP*), Dummy (*D*), Mimed Dummy (*MD*), Smeared Plant (*S*), Soil Infused (*INF*), and Predator Removal (*PR*); Risk Absence (*RA*) was the model reference. Significant results are shown (bold). See the main text Methods for experiment set-up and (Supplementary Fig. S1) for an illustration of off-plant aphid proportions, Resid. D.f. (residual degrees of freedom).

| Explanatory variables | Aphid proportions off plant | | | |
|---|---|---|---|---|
| | LR$\chi^2$ | Df | *Resid. D.f.* | P |
| Aphid density | 0.2 | 1 | 109 | 0.65 |
| Risk Treatment | 32.62 | 10 | 109 | **0.0003** |
| Aphid density × Risk Treatment | 6.06 | 10 | 109 | **0.81** |
| Aphid density × Risk Treatment × Census Day | 30.65 | 11 | 109 | **0.001** |

(Dispersion parameter for quasipoisson family taken to be 1.98193)
Null deviance: 320.20 on 141 degrees of freedom
Residual deviance: 182.47 on 109 degrees of freedom

**Fig. S1. Aphid proportions off plant under predator-associated risk treatments.** Average proportions of green peach aphid *M. persicae* that were found off host plants in the microcosm, upon two censuses [Day 4 and Day 8]), in response to the risk treatments associated with ladybird *C. septempunctata* comprising 11 levels: Live Predator (*LP*), Oscillated Live Predator (*OLP*), Isolation I (*ISO1*), Isolation II (*ISO2*), Dead Predator (*DP*), Dummy (*D*), Mimed Dummy (*MD*), Smeared Plant (*S*), Soil Infused (*INF*), and Predator Removal (*PR*); Risk Absence (*RA*) was the model reference. See the main text Methods for experiment set-up. The orange circles show the off-plant proportions on Day 4 (1st census); the blue circles show off-plant proportions on the final day of the experiments (2nd census).

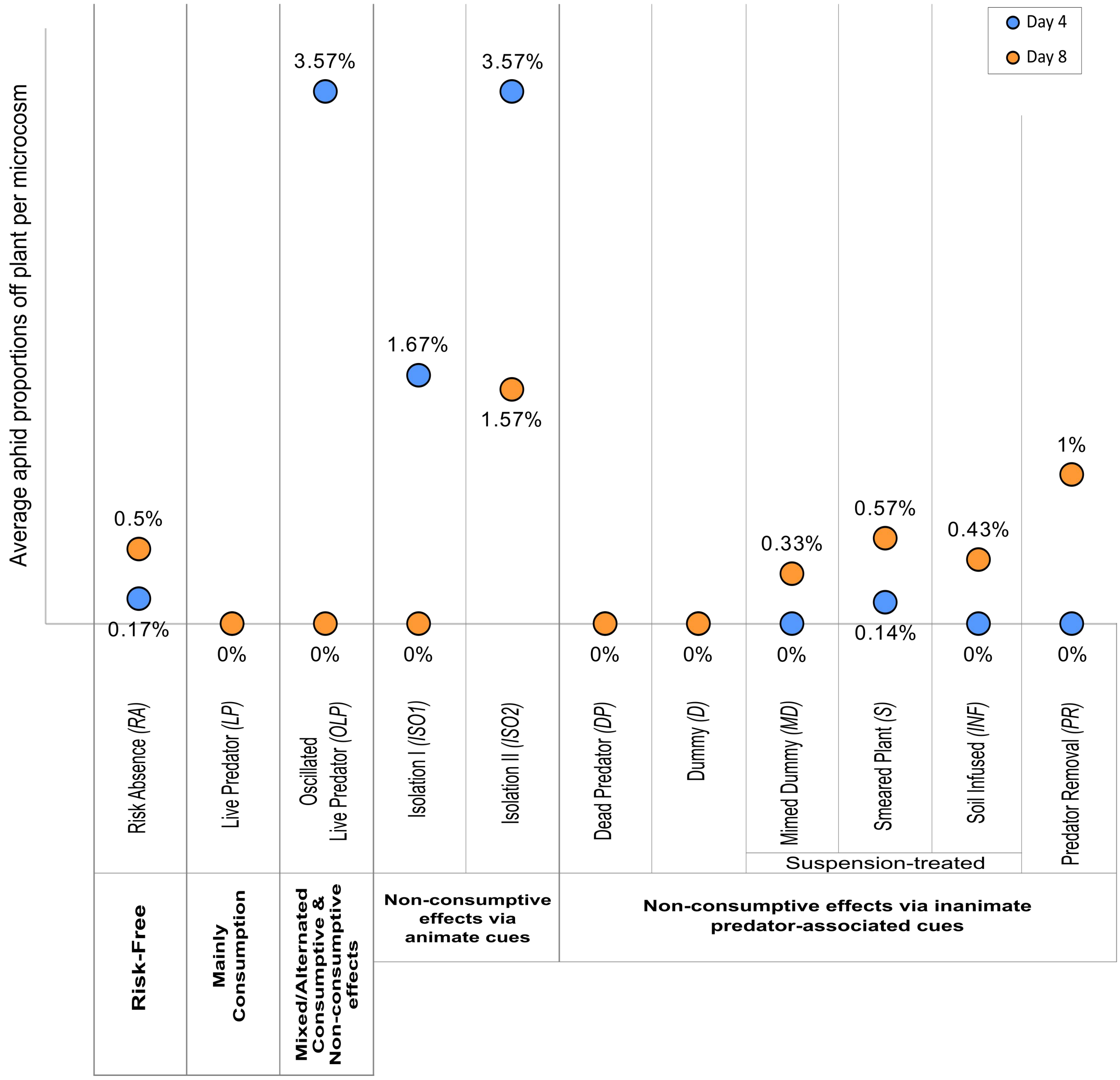
